# JEDII: Juxtaposition Enabled DNA-binding Interface Identifier

**DOI:** 10.1101/2022.05.19.492702

**Authors:** Sanjana Nair, M.S. Madhusudhan

## Abstract

The specific recognition of small stretches of the genomic sequence by their cognate binding protein partners is crucial for various biological processes. Traditionally the prediction of DNA-protein interactions has been treated as two separate problems - one where we predict the most probable DNA sequence that a given protein would bind to and another where we determine the amino acids constituting the DNA binding pocket on a protein. In this study, we introduce JEDII, a template-based method that combines these two aspects of DNA-protein interactions and predicts the residues, nucleotides and amino acids, that would mediate the interaction. Our computational method utilises known structures of DNA-protein complexes in a protocol that superimposes amino acid-nucleotide hydrogen-bonding donor and acceptors atoms on one another to identify the protein-DNA interface. The corner stone of the method is that specificity bestowing hydrogen-bonding interactions are structurally conserved. We validated the accuracy of our procedure on a dataset of 285 DNA-protein complexes where JEDII predicted the cognate DNA sequence with a 62% accuracy. It predicted the DNA-binding amino acids on the protein with 94 % accuracy and an MCC of 0.70. JEDII was also separately compared to other popular methods that predict the cognate DNA sequence and to methods that predict the DNA binding residues. The comparisons were done over four different datasets and JEDII outperformed most methods over all these data sets. JEDII is a robust method following a simple replicable algorithm to determine the molecular basis of DNA-protein specificity and could be instrumental in predicting DNA-protein complexes that are central to key biological phenomena.

## Introduction

Many crucial cellular processes such as DNA replication (1, 2), DNA-repair (3, 4), transcription (5–7), regulation of gene expression (8, 9), cellular differentiation (10–13), genome organization (14), etc. depend on the interaction of protein and DNA. The three-dimensional (3D) structures of the protein-DNA complexes resulting from these interactions provide mechanistic details of their function (15–20). These structural data can provide information at the level of individual residues. This information from structures could help identify/rationalize the residues at the DNA-protein interface whose disruption could lead to diseased conditions (21–23).

The experimental determination of DNA-protein structures is time-consuming, and the structures of all protein families are not represented. Even among the protein families with DNA-protein complex data, the sampling of the constituent members is not uniform in the Protein Data Bank (PDB) (24). There are also experimental methods that provide indirect information about the DNA-protein binding interface, such as-extensive mutagenesis combined with Electrophoretic Mobility Shift Assay (EMSA) (25, 26) and crosslinking-coupled with mass spectroscopy (27, 28), Chromatin Immuno Precipitation Sequencing (ChIP-Seq) (29–32), protein binding microarrays (33, 34), DNase I footprinting (35) and Systematic Evolution of Ligands by EXponential enrichment (SELEX) (36). However, the slow progress by experimental methods in generating DNA-protein interface data initiated the development of a plethora of computational methods. These methods can provide high throughput results in a relatively short time.

The prediction of the DNA-protein interface has been treated as two separate problems. One set of methods solely predicts the DNA-binding residues, while a second set looks at predicting the DNA sequence recognized by a protein.

Among the methods that predict DNA-binding residues, there are three classes based on the features they use to make the prediction-sequence-based, structure-based and a combination of sequence and structural features.

Sequence-based methods rely only on the protein sequence to predict the binding residues. The sequence-based algorithms employ various methods for predicting DNA-binding residues such as Support Vector Machines (SVMs) (37–44). neural networks (45–50), random forest classification (51, 52), ensemble learning (53), deep learning (54) and other combinations of machine learning techniques (55–63).

Structure-based methods exclusively employ features from structural data to predict DNA-binding residues such as electrostatics, geometry and surface curvature (64–68).

The final class of methods uses sequence and structural information for DNA-binding residue prediction. Since they incorporate all available data for a given protein, they are relatively more accurate than sequence or structure-based methods. The methods that use both sequential and structural features are more diverse in the techniques that they use for predicting DNA-binding residues. The techniques include SVM (43, 69–71), ensemble learning (72), neural networks (73–75) docking (76), clustering (77–79), light gradient boosting (80), and a combination of machine learning and template-based techniques (81).

The above-mentioned methods have successfully predicted the DNA-binding residues to various degrees. However, predicting DNA-protein interface not only involves predicting amino acid interacting partners, but also necessitates predicting the DNA sequence for a protein. Most of the computational methods for the prediction of DNA sequence deal with transcription factor binding sites (82, 83, 92, 84–91). These methods require a DNA sequence as an input and search for the transcription factor binding sites within this region. However, there are also methods that predict the DNA sequence for all DNA-binding proteins with DNA sequence alone or DNA and protein sequences as input (93). Some prediction tools also use structural data as input (94–97). However, no single method identifies both DNA and protein binding regions.

Our proposed method does not treat the identification of DNA-protein interface identification as separate problems and predicts the binding residues on protein as well as DNA. The method employs topology independent structural superimpositions to derive the DNA-binding interface. Ours is a template-dependent algorithm that relies on the hydrogen bond donor-acceptor pattern similarity at the DNA-protein interface to identify appropriate template structures. We deal with the problem of DNA-binding interface prediction on the whole. Therefore, our method predicts the DNA-binding residues, binding pose, and the putative DNA sequence for a given protein structure. To evaluate the efficacy of our method, we employed a dataset curated by us and compared ourselves to four other datasets (45, 68, 74, 78). We evaluated the robustness of the method in the absence of close homologs, with unbound protein conformations and protein models as inputs. We validated the DNA sequence prediction for a query protein, with ChIP seq data and compared the efficacy with DBD2BS (97) and the Farrel and Guo method (96).

## Methods

### 1. Dataset Curation

The datasets used for the prediction of the DNA-binding interface (algorithm described in section 2) are -

#### 1a) Query Dataset

We followed the steps below to construct the query dataset -

i. We obtained 3467 DNA-protein complex structures from RCSB PDB (98) by using the following parameters - ‘Molecule Type=complex, Experimental Method=X-RAY, and Chain Type: there is a Protein, and a DNA chain but not any RNA or Hybrid and Resolution is between 0.0 and 3.0 Å.’ (April 2018).
ii. To avoid over-representation of protein families, we compiled a 40% non-redundant set using h-CD-HIT. Multiple iterations result in more efficient and more accurate clustering (99). Therefore, we performed three iterations at 90,60 and 40% identity, which resulted in 669 clusters.
iii. We identified DNA-protein hydrogen bonds for each structure in all the 669 clusters using an in-house python script. The maximum distance between donor and acceptor atoms was 4Å, and the angle between donor-acceptor-acceptor antecedent was between 90-180° (100). We chose a permissive (larger) distance cutoff and angle range to include all potential hydrogen-bonded amino acid-nucleotide pairs. The cluster representative was the structure with the maximum number of DNA-protein hydrogen bonds in a chain. Wherever there were multiple chains with maximum hydrogen bonds, the structure with the higher total number of hydrogen bonds (all chains included) was chosen. If two or more structures had the same number of overall hydrogen bonds, the structure with the better resolution was the representative. From 669 clusters, we removed 81 clusters with no DNA-protein hydrogen bonds, which left us with 588 clusters with 588 representative structures.
iv. In this study we only considered the classical conformation of DNA (B-form double helix). Thus, we removed 166 structures after screening for double-stranded B-form DNA using Nucleic Acid Database (NDB), resulting in 422 structures (101). NDB missed filtering out some entries that did not belong to the double-stranded B-form DNA category. We wanted to include only the DNA-protein complexes that exhibited DNA-sequence-specific binding since our method also predicts the DNA sequence recoognized by the protein. Therefore, we did keep structures that had non-specific DNA-binding activity, such as polymerases. We manually curated the entries to remove both these types of structures by text mining and visualization of the DNA-protein complexes. The final dataset consisted of 285 structures, which we termed the PDNA-285 dataset. Four of the structures in the PDNA-285 dataset contained heteromeric proteins interacting with DNA, and therefore the dataset has 289 unique proteins. In homo-oligomeric protein assemblies, we retained unique chains if they had non-overlapping binding sites (The PDB IDs and chains used in the PDNA-285 dataset are mentioned in Supplementary Table 1).

#### 1b) Template dataset

The template database consisted of only X-ray crystal structures having a resolution better than 3Å and composed only of DNA and protein chains that resulted in 3909 structures (May 2020). Wherever rotational data was present, we performed symmetry operations and used the biological unit of these structures. We obtained a 40% NR template database with 700 sequences using these 3909 PDBs as input to h-CD-HIT to assess the performance of our method in the absence of close homologs in the template dataset.

#### 1c) Benchmark datasets

We validated the prediction of amino acids binding DNA using datasets from previous studies-PDNA-62 (45) dataset, PDNA224 (71),, PDNA-129 (74), HOLO-APO 82 (78), where the numbers in the dataset name indicate their size.

For the validation of DNA-sequence prediction, apart from the PDNA-285 dataset, we have used three datasets-

i. A set of 7 proteins used by DBD2BS in their study.
ii. Position Weight Matrices (PWMs) generated by ChIP seq data from JASPAR for comparison with the predicted DNA sequence (102). JASPAR had 331 entries for PWMs, of which 72 had a three-dimensional (3-D) protein structure. These proteins did not possess a 3-D structure in complex with the DNA and formed the validation set for DNA sequence prediction.
iii. 27 Transcription factor chains from the Farrel and Guo study (96): Their dataset had 27 protein chains, which bound as monomers to their binding site. We obtained the PWMs for each of these proteins from JASPAR. We selected the nucleotide with highest frequency at each position to generate reference DNA sequences against which we evaluated our method.

#### 1d) Validation using models

All previously described validations used crystal structures extracted from the PDB. We wanted to evaluate the efficacy of our prediction method when using models of of proteins instead of their experimentally determined structures. To this end, we constructed models of the proteins in the PDNA-129 dataset separately using MODELLER (103) and AlphaFold2 (104).

For generating models with MODELLER, we used the protein with the best sequence identity with the query excluding the crystal structure as the template for modelling. We obtained 122 models as seven sequences did not find alternate templates. We analyzed the TM-score of these 122 models using TM-align to ascertain their similarity to the experimental structures (105). 97 of the 122 models had a TM-score >0.5, where a higher TM-score indicates greater structural similarity.

We also modeled the sequences from PDNA-129 using AlphaFold2 (using either the AlphaFold2 database or modelling with the google colab notebook). 127 of these 129 sequences have a TM-score >0.5.

### 2. Prediction Algorithm

The prediction algorithm relies on the similarity of the hydrogen bond pattern at the interface of DNA-protein interactions. We used hydrogen bond donor and acceptor atoms in protein residues to guide structural alignments using CLICK (106, 107). The hydroxyl group in serine, threonine and tyrosine can act as both donor and acceptor. As a practical consideration, we treated this group as a donor in this study, since it acts as a donor in 70% of cases in the PDNA-285 dataset. We used default CLICK settings to align the query structure with all structures in the template database (except for the same PDB structure). The default settings use solvent accessibility and secondary structure parameters for alignment.

The DNA coordinates of the template were transferred to the superimposed query to build a model. To remove unsatisfactory models, we used the criteria of clashes and contacts. A *clash* was defined as a distance less than 1.5 Å between the C, C^α^ atom of the protein, and the DNA atoms. We discarded all models with more than one clash.

The O and N atoms of the main chain are not explicitly considered in the checking of clashes as these atoms are hydrogen bond acceptors and donors, respectively. Our method involves the superimposition of the query with a template and the subsequent transfer of coordinates of the DNA to form the DNA-protein complex. This complex is not energy minimized or processed in any other way after superimposition. The clash criteria is the only means of eliminating possible bad models. This is also why the clash criteria is somewhat permissive (1.5 Å threshold, instead of a larger value). In addition, we have also made the hydrogen bond criteria permissive as we are allowing all donor-acceptor distances below 4 Å. Potentially these could include clashes (below 1.5 Å) - but we observed that this occurs in ~1% of the N and O atoms in the data (Appendix 19). With these permissive hydrogen bond and clash criteria, we believe we can identify all plausible hydrogen-bonded pairs.

A *contact* between DNA and the protein consisted of a distance less than 3.5 Å between the hydrogen bond donor/acceptor atoms of protein and DNA. We removed the models with less than three contacts between DNA and protein.

To select the best model, we define a term called the query overlap percentage -

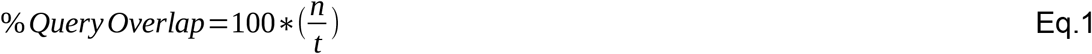

where *n* is the number of donors and acceptors in the query protein superimposed with the template and *t* is the total number of donors and acceptors in the query protein. We predicted the model with the highest query overlap as the putative binding pose. The DNA-binding residues were the amino acid residues with any atoms within a distance of 4 Å from any DNA atom. We predicted the DNA sequence of the template as the putative nucleotide sequence that the protein would bind. We labeled this protocol of predicting the DNA-binding interface as Juxtaposition Enabled DNA-binding Interface Identifier (JEDII) (Figure 1). We demonstrated the efficacy of JEDII using PDNA-285 dataset and validated it using the different benchmark datasets.

**Figure 1:**
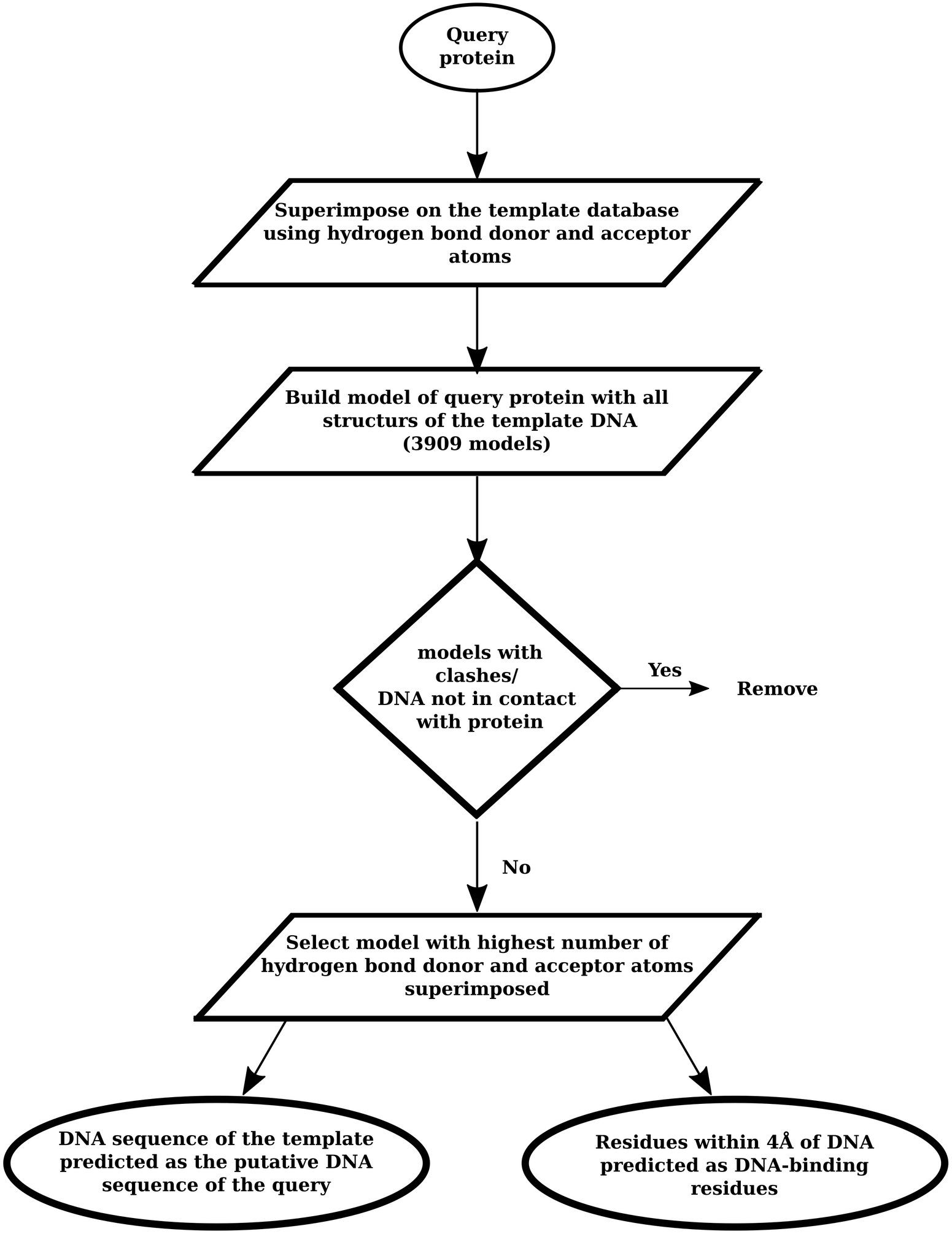
Flowchart of the steps in the JEDII protocol

### 3 Comparison of JEDII with GraphBind and HDock on PDNA-285 dataset

We compared the performance of JEDII with GraphBind (74) and HDock (108, 109) on the PDNA-285 dataset. We used the GraphBind standalone version to make predictions on proteins from the PDNA-285 dataset. Out of 285 PDBs in our dataset, 107 are a part of the GraphBind training set. Therefore, we predicted the DNA-binding residues for the remaining 178 proteins. We parallelized GraphBind jobs using GNU Parallel (110). For predictions using HDock, we docked the DNA structure from PDB ID 6EL8 on each of the proteins in the dataset and predicted the residues within 4 Å of the DNA in the top model as the DNA-binding residues.

### 4 Evaluation Metrics

We evaluated the predictions of amino acid residues binding DNA using Matthew’s Correlation Coefficient (111) (MCC), Specificity (SP), Sensitivity (SN), Precision (P), Accuracy (Acc) and F1 score. We have reported values in % for SP, SN, P and Acc.

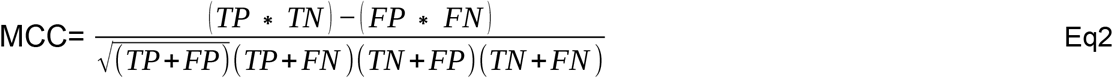

Where,

*TP* = true positive (predicted binding residue correctly),
*TN*= true negative (predicted non-binding residue correctly),
*FP*= false positive (predicted a non-binding residue as a binding residue),
*FN*= false negative (predicted a binding residue as non-binding).

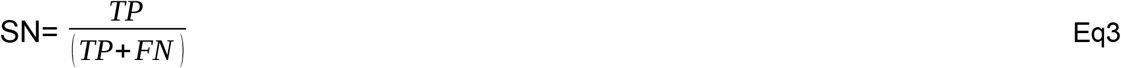

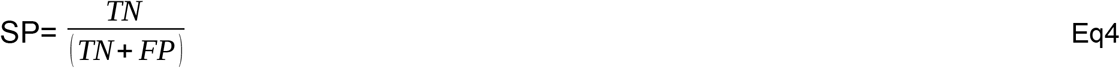

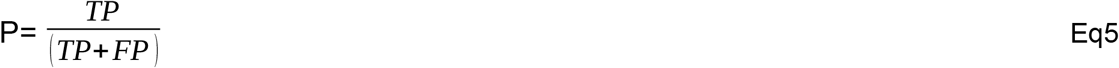

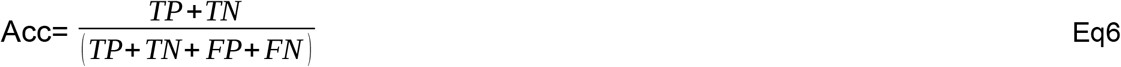

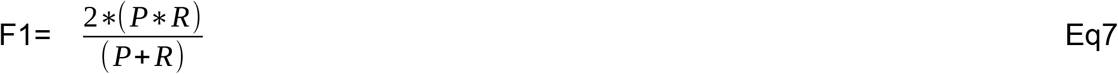

In the JEDII protocol, we selected the template DNA sequence as the sequence of the DNA likely to bind the input protein. To evaluate the prediction, we aligned the predicted sequence and the actual sequence using the malign module of MODELLER. We then calculated the similarity score as -

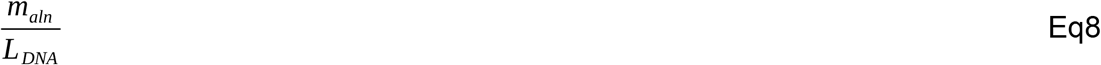

where *m_aln_* is the number of matched nucleotides and *L_DNA_* is the length of the actual DNA sequence.

## Results

### 1. Assessment of the robustness of JEDII in predicting the DNA-protein interface

#### A) Performance on the PDNA-285 dataset

We predicted the DNA-binding interface for the proteins in the PDNA-285 dataset. We have segregated the prediction of the DNA-protein interface into the predictions of amino acid residues binding DNA and the prediction of DNA sequence that would bind the protein. In each section, we have described the results for predicting amino acid residues binding DNA, followed by the DNA sequence prediction.

From the predicted DNA-protein model for each query protein, we annotated all the amino acid residues within 4Å of the DNA as the DNA-binding residues. Using this criterion, JEDII achieved an overall accuracy of 94% and an MCC of 0.70 on this dataset (Table 1). The MCC for the dataset shows a bimodal distribution with one maxima at 0 and another at 0.9 (Figure 2 (a)).

**Figure 2:**
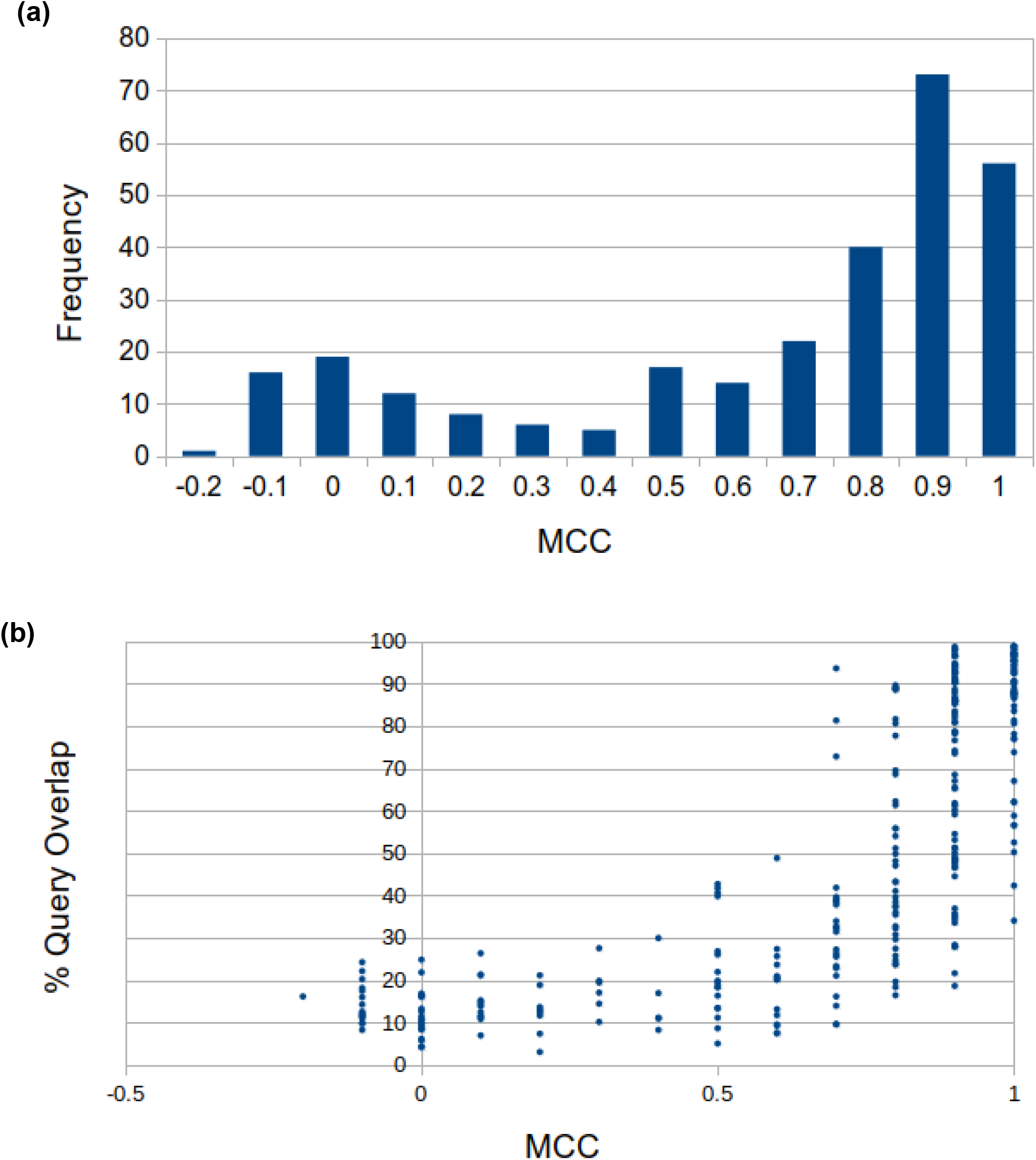
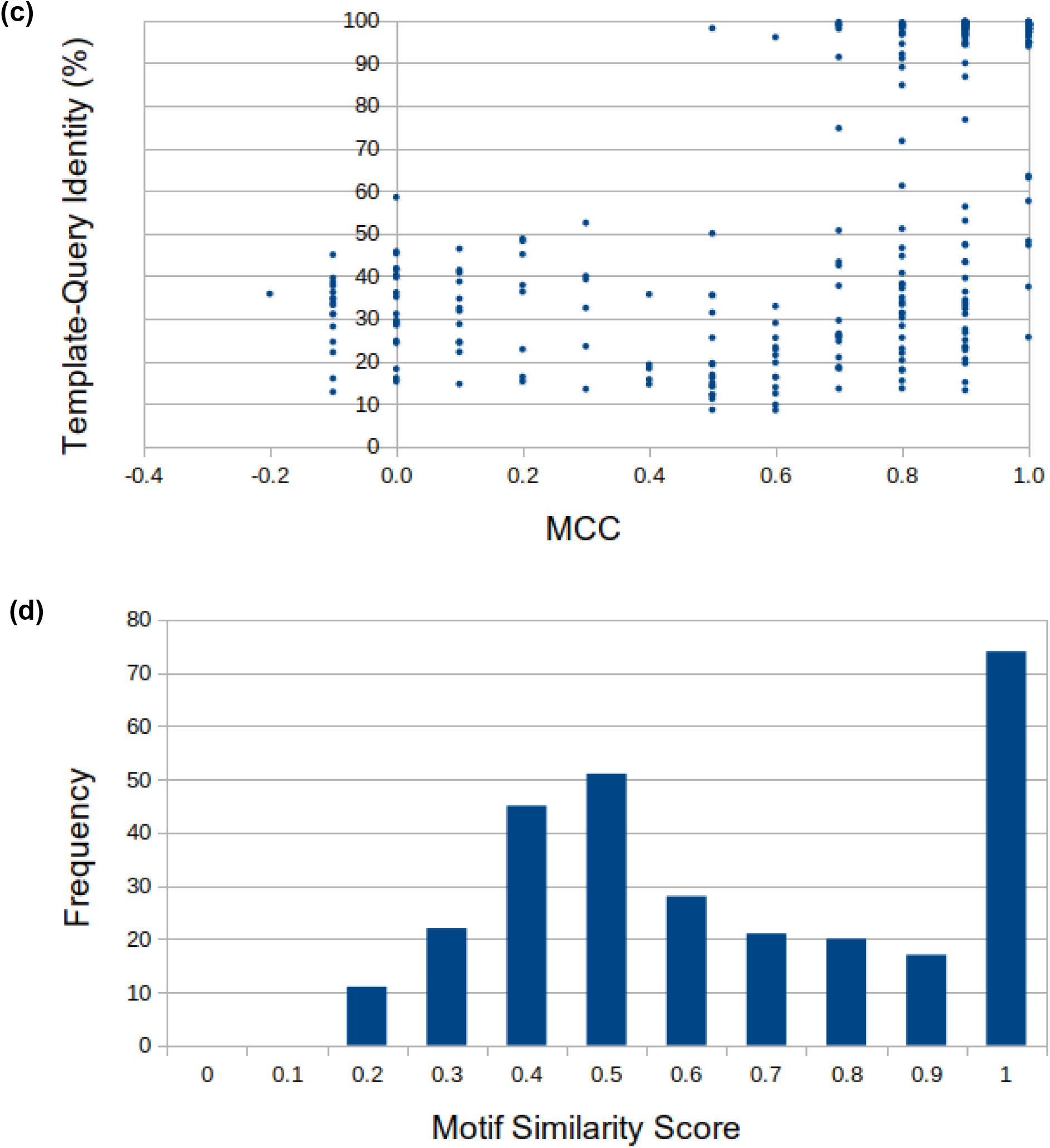
Performance of JEDII on the PDNA-285 dataset Subplots (a), (b) and (c) show MCC distribution, variation in MCC with query overlap (%) and average sequence identity between template-query (%), respectively for the prediction of DNA-binding residues. (d) is the distribution of similarity score for the DNA-sequence prediction.

**Table 1:** Performance of JEDII on PDNA-285 dataset with redundant and 40% non-redundant template database. The terms Acc, MCC, SN, P and F1 stand for Accuracy, Matthew’s Correlation Coefficient, Sensitivity, Precision and F1 score, respectively.

| Method | Dataset | Acc (%) | MCC | SN (%) | SP (%) | P (%) | F1 |
| --- | --- | --- | --- | --- | --- | --- | --- |
| JEDII | Redundant | 94 | 0.70 | 65 | 98 | 84 | 0.73 |
| JEDII | NR-40 | 90 | 0.45 | 38 | 97 | 66 | 0.48 |

We evaluated two factors contributing to the performance of the methodquery overlap and sequence identity between the template and the query. We plotted these parameters in the PDNA-285 dataset against the MCC (Figure 2(b) and 2(c)). The average query overlap was 49.1± 32.5%, while the average sequence identity between template-query was 59.1 ± 34.6 %. The query overlap exhibits a stronger positive correlation with the MCC (Spearman’s correlation Coefficient= 0.84) than the sequence identity between the template and the query (Spearman’s correlation Coefficient=0.60).

87.5% of the perfectly-identified binding sites (49 out of 56) have a template identity >90%. Among the remaining seven proteins with a DNA-binding residue prediction of 1, the lowest sequence identity between the template and query was 25.6% (PDB ID: 1R8D). Moreover, in 14 PDBs with MCC ≥ 0.4, the template selected had a different CATH (112, 113) fold from the query.

JEDII had an average similarity score of 0.62 for predicting the DNA sequence on the PDNA-285 dataset (Figure 2 (d)). For 62 out of 289 unique proteins, we could identify the DNA sequence with a similarity score of 1.

An example of the identifying the correct DNA-protein interface with a sequentially and structurally different template is that of the cro repressor.

We predicted the DNA-binding pose and the DNA sequence for cro repressor of bacteriophage lambda (PDB ID: 6CRO). The template selected by JEDII was cI protein of bacteriophage lambda (PDB ID: 1RIO), which had a sequence identity of 31% with the cro repressor. The two proteins have different CATH IDs – 3.30.240.10 (6CRO), and 1.10.260.40,1.10.10.10 (1RIO) and the superimposition occurs between atoms in an α helix (cl) and atoms in a β sheet (cro repressor) (Figure 3). The predicted and the actual binding pose have the angle between the helical axis as 18.22° and the RMSD as 0.78Å. The MCC for the DNA-binding residue prediction is 0.9, and when compared with the DNA sequence in the experimental structure, the predicted DNA sequence has 14/16 nucleotides matched (0.87 similarity score).

**Figure 3:**
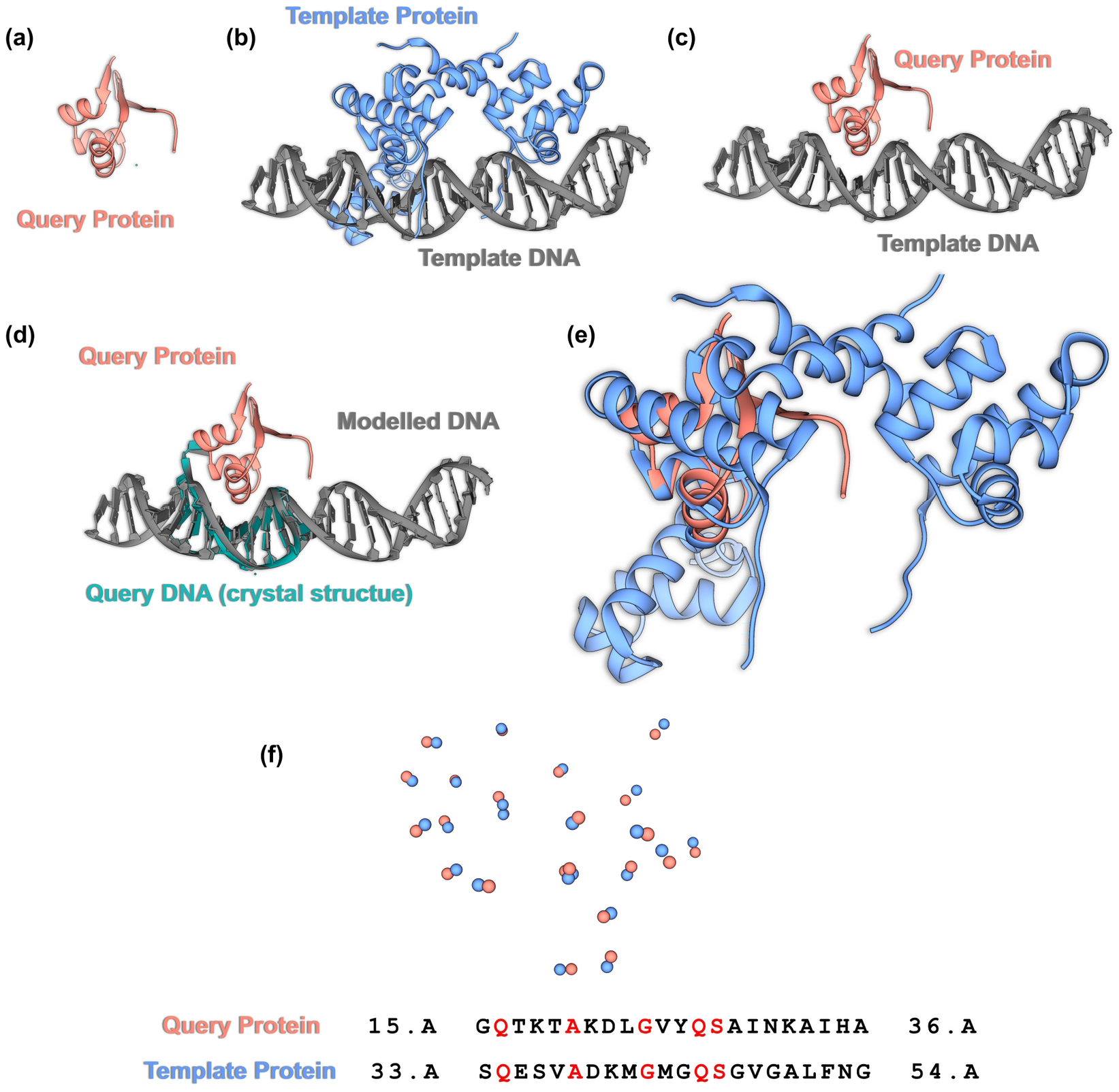
The prediction of the binding pose of cro repressor. The query and template proteins are in salmon and blue color, respectively. The query and template DNA are depicted in green and grey, respectively. (a) Structure of the query protein. (b) Structure of the template DNA-protein complex. (c) Structure of the predicted model. (d) Relative orientation of the DNA in the model and crystal structure of the query protein with DNA. (e) Superimposition of the query and template proteins using hydrogen bond donors and acceptors. (f) A section of the hydrogen bond donor/acceptor atoms of query (salmon) and template (blue) protein superimposed and the corresponding alignment of proteins. The amino acid residues with the same identity are highlighted in red.

#### B) Predictions in the absence of close homologs

We have established in A) that the accuracy of predictions is correlated with the query overlap (0.84) and the sequence identity (0.60) between template-query. However, sequential similarity also implies structural similarity, resulting in a better overlap of the query protein. Therefore, we wanted to assess the method’s robustness in the absence of close sequential homologs. We used 700 protein-DNA complexes from the NR-40 database as templates to make predictions for proteins in the PDNA-285 dataset. The accuracy of the method for DNA-binding residue prediction reduces to 90.0%, with an MCC of 0.45 (Table 1). We can still identify DNA-binding residues of 4 query proteins with an MCC of 1. The correlation between the MCC and the template-query sequence identity is −0.05, while that between MCC and % query overlap is 0.67. Therefore, in the absence of close homologs, the identification of DNA-binding residues relies on the structural similarity of the DNA-binding interface.

Without close sequential homologs in the template dataset, the prediction of the DNA sequence has an average similarity score of 0.40. Though there has been a reduction in the accuracy, JEDII could identify the DNA sequence for one of the query proteins (PDB ID: 5GZB) with a similarity score of 1.

#### C) Predictions with the unbound conformation of protein

We employed the HOLO-APO 82 dataset to estimate the efficiency of our method when provided with the unbound conformation of DNA. This dataset consists of 82 pairs of proteins in both apo (DNA-unbound) and holo (DNA-bound) conformation. The accuracy of the method for DNA-binding residue prediction is 95% and 93% for holo and apo conformation, respectively. The MCC of JEDII on the holo conformation is 0.67, while for the apo conformation, it is 0.55 (Table 5). In only 17 out of 82 cases, JEDII identified the corresponding holo-structure as the template. The average RMSD between apo and holo-structures was 1.2Å irrespective of whether JEDII identified the holo-structure as a template or not.

We studied the effect of the degree of conformational change on the efficacy of the method using correlations. We correlated the RMSD between the apo and holo-structures with the MCC of amino acid residues binding DNA. The Pearson’s correlation Coefficient for these two parameters was −0.08, indicating that the MCC is independent of the degree of conformational change. The better MCC for holo-structures could be due to their similarity with the structures of DNA-protein complexes in the template database.

JEDII had an average similarity score of 0.66 for predicting the DNA sequence on both the apo and holo datasets. Out of 82 sequences, the crystal structure DNA sequence was precisely identified (similarity score = 1) in 23 and 30 cases in holo and apo datasets, respectively.

#### D) Dependence of the accuracy on the use of models

We assessed the ability of our method to identify DNA-binding interface using models generated by MODELLER and AlphaFold2 from the protein sequences in the PDNA-129 dataset. The MCC values for the prediction of amino acids binding DNA on models built by MODELLER and AlphaFold2 are 0.32 and 0.33. Thus JEDII has an equivalent performance on both MODELLER and AlphFold2 models (with AlphaFold2 having a marginally better performance).

In the case of DNA sequence prediction with models as inputs, JEDII has a similarity score of 0.41 on MODELLER models and 0.40 on AlphaFold2 models. Perfectly identified DNA sequences (similarity score =1) are 6 and 7 for MODELLER and AlphaFold2 models, respectively.

### 2. Comparison of results with other methods on benchmark datasets

We wanted to assess the accuracy of our method in comparison to other existing algorithms. However, no software performs both functions of predicting amino acids binding DNA and DNA-sequence prediction. Therefore we compared JEDII with the different classes of methods separately.

#### A) Predicting amino acid residues binding DNA

##### A1) Comparison with other methods on benchmark datasets

We employed PDNA-62, P224 and PDNA-129 datasets that have been used in previous studies to analyze the performance of JEDII on DNA-binding residue prediction. We also compared the accuracy of JEDII with other methods on the APO-HOLO82 dataset and on models built from sequences derived from PDNA-129 dataset.

We have compared JEDII to sequence-based methods (ADASYN, BindN, BindN-RF, BindN+, CNNsite, DBS-Pred, DBS-PSSM, DNAPred, DRNAPred, EL_PSSM-RT, SVMnuc, TargetDNA, TargetS), structure-based methods (COACH-D) and methods that use a combination of both sequence and structural features (DNABind, DP-Bind, GraphBind, NucBind, PDNAsite, PDRLGB, PreDNA, RBSCORE). We obtained the values for all evaluation metrics from the respective papers for all methods except GraphBind. In case of GraphBind, the program was run on models from PDNA-129 and on the 178 proteins from PDNA-285 dataset.

On both the PDNA-62 and P224, we perform better than all other methods in every assessment measure except sensitivity (Table 2 and Table 3). On the PDNA-129 dataset, we perform better than or at par with most methods, except GraphBind. GraphBind performs better than JEDII in all measures except precision (Table 4). The PDNA-129 dataset contains 27 entries that have been solved using electron microscopy. Since our template database only has DNA-protein complexes with resolution <4 Å, we have a worse performance on this dataset compared to GraphBind.

**Table 2:**
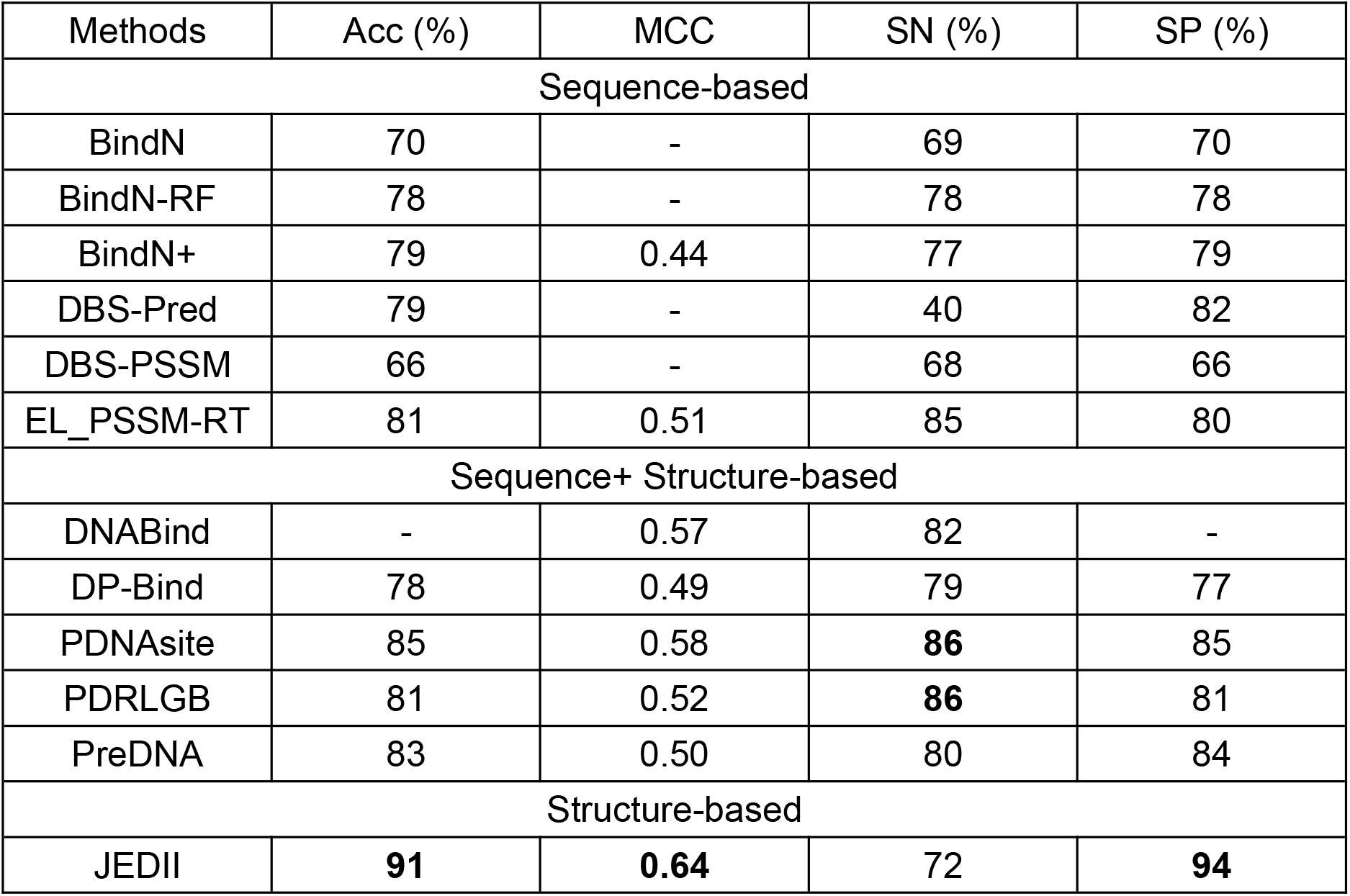
Comparing JEDII with existing algorithms on PDNA-62 Dataset. Values for BindN (37), BindN-RF (51), BindN+ (38), DBS-Pred (45), DBS-PSSM (46), EL_PSSM-RT (53), DNABind (81), DP-Bind (56), PDNAsite (72), PDRLGB (80) and PreDNA (71) were taken from their respective papers. The terms Acc, MCC, SN and SP stand for Accuracy, Matthew’s Correlation Coefficient, Sensitivity and Specificity, respectively.

**Table 3:** Comparing JEDII with existing algorithms on the P224 Dataset. The values for ADASYN (49), CNNsite (50), EL_PSSM-RT (53), PDNAsite (72), PDRLGB (80), PreDNA (71) and RBSCORE (79) were obtained from their respective papers. The terms Acc, MCC, SN and SP stand for Accuracy, Matthew’s Correlation Coefficient, Sensitivity and Specificity, respectively.

| Method | Acc (%) | MCC | SN (%) | SP (%) |
| --- | --- | --- | --- | --- |
| Sequence-based |  |  |  |  |
| ADASYN | 83 | 0.48 | 77 | 84 |
| CNNsite | 84 | 0.40 | 77 | 84 |
| EL_PSSM-RT | 78 | 0.34 | 80 | 78 |
| Sequence + Structure-based |  |  |  |  |
| PDNAsite | 82 | 0.40 | **83** | 82 |
| PDRLGB | 80 | 0.38 | **83** | 80 |
| PreDNA | 82 | 0.35 | 76 | 82 |
| RBSCORE | 85 | 0.40 | 63 | 88 |
| Structure-based |  |  |  |  |
| JEDII | **94** | **0.66** | 68 | **97** |

**Table 4:** Comparing JEDII with existing algorithms on the PDNA-129 dataset. Values for DNAPred (39), SVMnuc (43), TargetDNA (44), TargetS (63), DNABind (81), GraphBind (74), NucBind (43) and COACH-D (64) were taken from their respective papers. The terms Acc, MCC, SN, P and F1 stand for Accuracy, Matthew’s Correlation Coefficient, Sensitivity, Precision and F1 score, respectively.

| Method | MCC | SN (%) | P (%) | F1 |
| --- | --- | --- | --- | --- |
| Sequence-based |  |  |  |  |
| DNAPred | 0.33 | 40 | 35 | 0.37 |
| SVMnuc | 0.30 | - | 37 | 0.34 |
| TargetDNA | 0.29 | 42 | 28 | 0.34 |
| TargetS | 0.26 | 24 | 37 | 0.29 |
| Sequence+ Structure-based |  |  |  |  |
| DNABind | 0.41 | 60 | 35 | 0.44 |
| GraphBind | **0.50** | **68** | 42 | **0.52** |
| NucBind | 0.31 | - | 37 | 0.35 |
| Structure-based |  |  |  |  |
| COACH-D | 0.30 | - | 36 | 0.34 |
| JEDII | 0.41 | 42 | **47** | 0.45 |

JEDII performs better on the HOLO-APO 82 dataset as compared to other methods. (Table 5). While on the holo dataset, JEDII outperforms other methods in all evaluation parameters, in the apo dataset, it has lower F1 and sensitivity compared to JET2DNA.

**Table 5:**
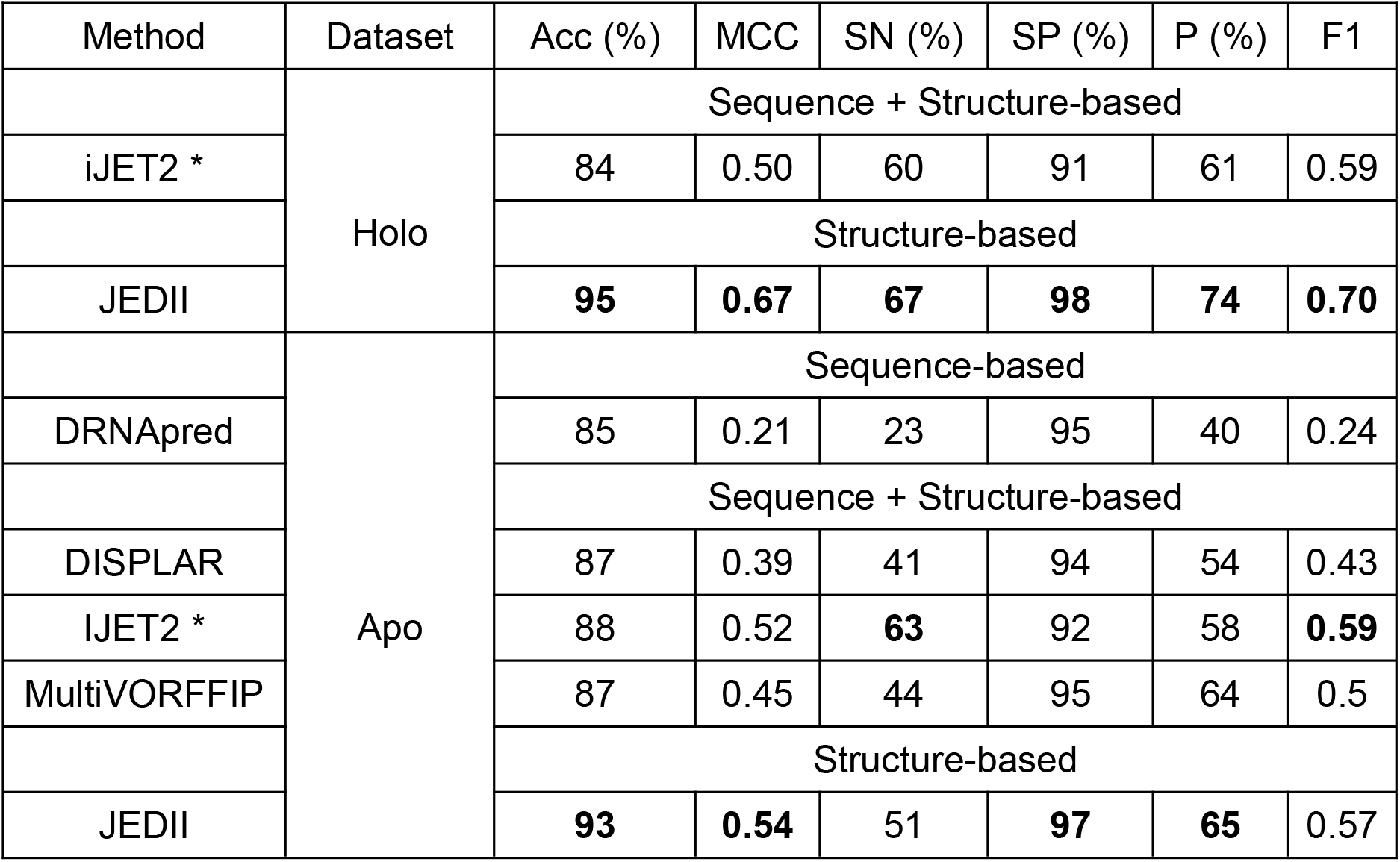
Comparing the performance of JEDII on the HOLO-APO 82 dataset with other methods. *iJET2 has divided the holo set to two parts (74 and 8 proteins) based on the stoichiometry. For comparison, we have used the average value of the evaluation metrics. Values for iJET2 (78), DRNAPred (58), DISPLAR (73) and MultiVORFFIP (76) were taken from their respective papers. The terms Acc, MCC, SN, SP, P and F1 stand for Accuracy, Matthew’s Correlation Coefficient, Sensitivity, Specificity, Precision and F1 score, respectively.

We compared the prediction of DNA-binding residues from protein models with GraphBind. We used the models built by MODELLER and AlphaFold2 for evaluation. On both, the models built by MODELLER and AlphaFold2, JEDII has a better accuracy, specificity and precision but GraphBind has better MCC and F1 values (Table 6). This trend holds even when we only consider the models that have TM-score >0.5.

**Table 6:**
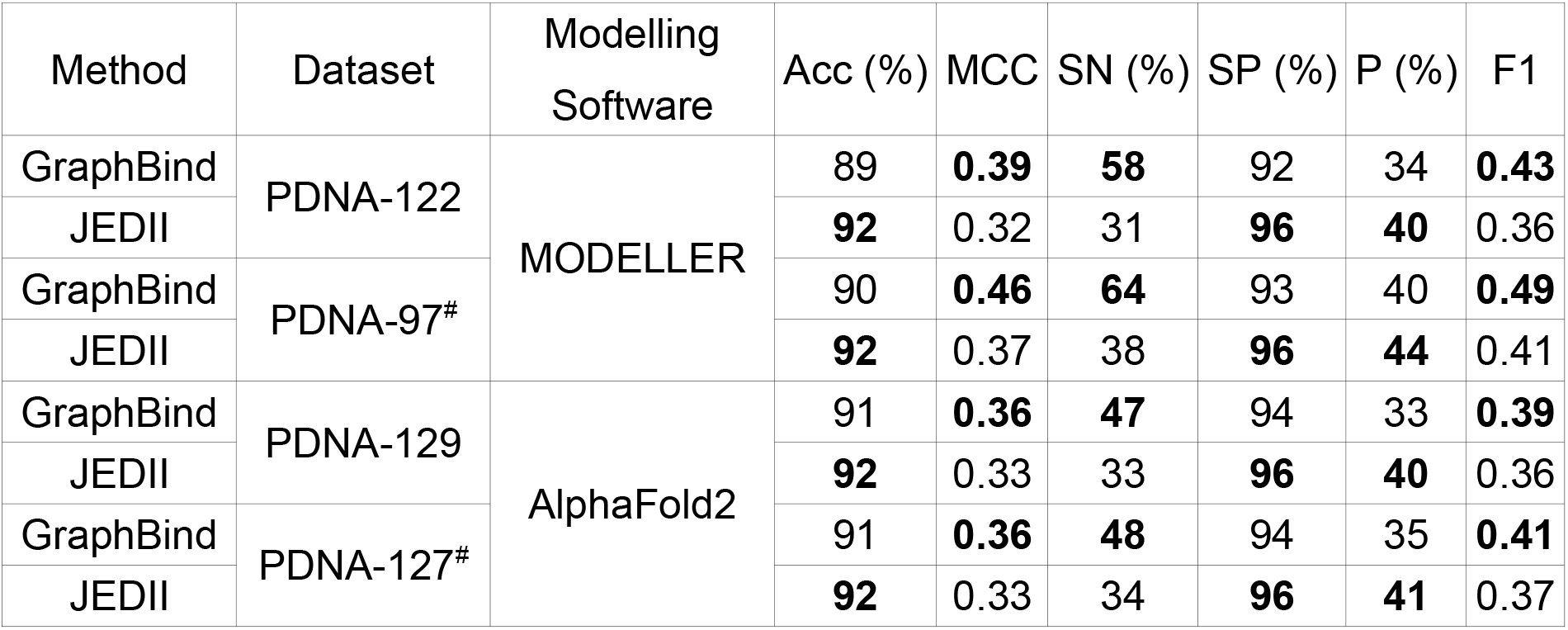
Comparing the accuracy of JEDII on models built by MODELLER and AlphaFold. # indicates models with TM-scores >0.5. The values for GraphBind were obtained by running the standalone version (74). The terms Acc, MCC, SN, SP, P and F1 stand for Accuracy, Matthew’s Correlation Coefficient, Sensitivity, Specificity. Precision and F1 score, respectively.

| Method | Dataset | Modelling Software | Acc (%) | MCC | SN (%) | SP (%) | P (%) | F1 |
| --- | --- | --- | --- | --- | --- | --- | --- | --- |
| GraphBind | PDNA-122 | MODELLER | 89 | **0.39** | **58** | 92 | 34 | **0.43** |
| JEDII |  |  | **92** | 0.32 | 31 | **96** | **40** | 0.36 |
| GraphBind | PDNA-97<sup>#</sup> |  | 90 | **0.46** | **64** | 93 | 40 | **0.49** |
| JEDII |  |  | **92** | 0.37 | 38 | **96** | **44** | 0.41 |
| GraphBind | PDNA-129 | AlphaFold2 | 91 | **0.36** | **47** | 94 | 33 | **0.39** |
| JEDII |  |  | **92** | 0.33 | 33 | **96** | **40** | 0.36 |
| GraphBind | PDNA-127<sup>#</sup> |  | 91 | **0.36** | **48** | 94 | 35 | **0.41** |
| JEDII |  |  | **92** | 0.33 | 34 | **96** | **41** | 0.37 |

##### A2) Comparison with other methods on the PDNA-285 dataset

For comparison of our accuracy, with other methods on the PDNA-285 dataset, we used GraphBind and HDock. JEDII performs better than GraphBind and HDock on the PDNA-285 dataset in all measures except sensitivity, where GraphBind is better (Table 7). Overall, except for PDNA-129, JEDII has a better performance than other sequence-based, structure-based and methods using both sequence and structure data. We consistently have a better specificity than other methods, however sensitivity of the method is one of the limitations of the method.

**Table 7:** Comparing JEDII with GraphBind and HDock on PDNA-285 dataset. * As mentioned in the methods, predictions were made for 178 proteins not in the GraphBind training set. The terms Acc, MCC, SN, SP, P and F1 stand for Accuracy, Matthew’s Correlation Coefficient, Sensitivity, Specificity. Precision and F1 score, respectively.

| Method | Dataset | Acc (%) | MCC | SN (%) | SP (%) | P (%) | F1 |
| --- | --- | --- | --- | --- | --- | --- | --- |
| GraphBind | PDNA-285* | 92 | 0.69 | **86** | 93 | 63 | 0.73 |
| HDock | PDNA-285 | 91 | 0.55 | 49 | 97 | 72 | 0.58 |
| JEDII | PDNA-285 | **94** | **0.70** | 65 | **98** | **84** | **0.73** |

**Table 8:** Number of nucleotide positions where the predicted nucleotide also had the highest frequency of occurrence in JEDII and DBD2BS.

| Sr. No | PDB ID | JEDII | DBD2BS |
| --- | --- | --- | --- |
| 1 | 1NFI | 5/10 | 5/10 |
| 2 | 1MI7 | 9/19 | 5/19 |
| 3 | 2A63 | 13/17 | 7/17 |
| 4 | 1M6U | 3/12 | 6/12 |
| 5 | 1GV2 | 4/11 | 5/11 |
| 6 | 1MH3 | 7/18 | 6/18 |
| 7 | 2GZW | 9/10 | 5/10 |

#### B) DNA sequence prediction

##### B1) Validation of DNA sequence prediction on ChIP seq data

ChIP seq experiments generate DNA-binding sequence data for proteins, that is generally stored as PWMs in databases such as JASPAR. We took 72 proteins from JASPAR that did not a structure in complex with DNA, but had a structure of the protein to assess the accuracy of JEDII in DNA sequence prediction. To derive the DNA-binding site, we expanded the PWMs to individual sequences that satisfy the PWM criteria. When we compared the DNA sequence predicted by JEDII with the sequences from PWM, we obtained an average similarity score was 0.81 (Figure 4).

**Figure 4:**
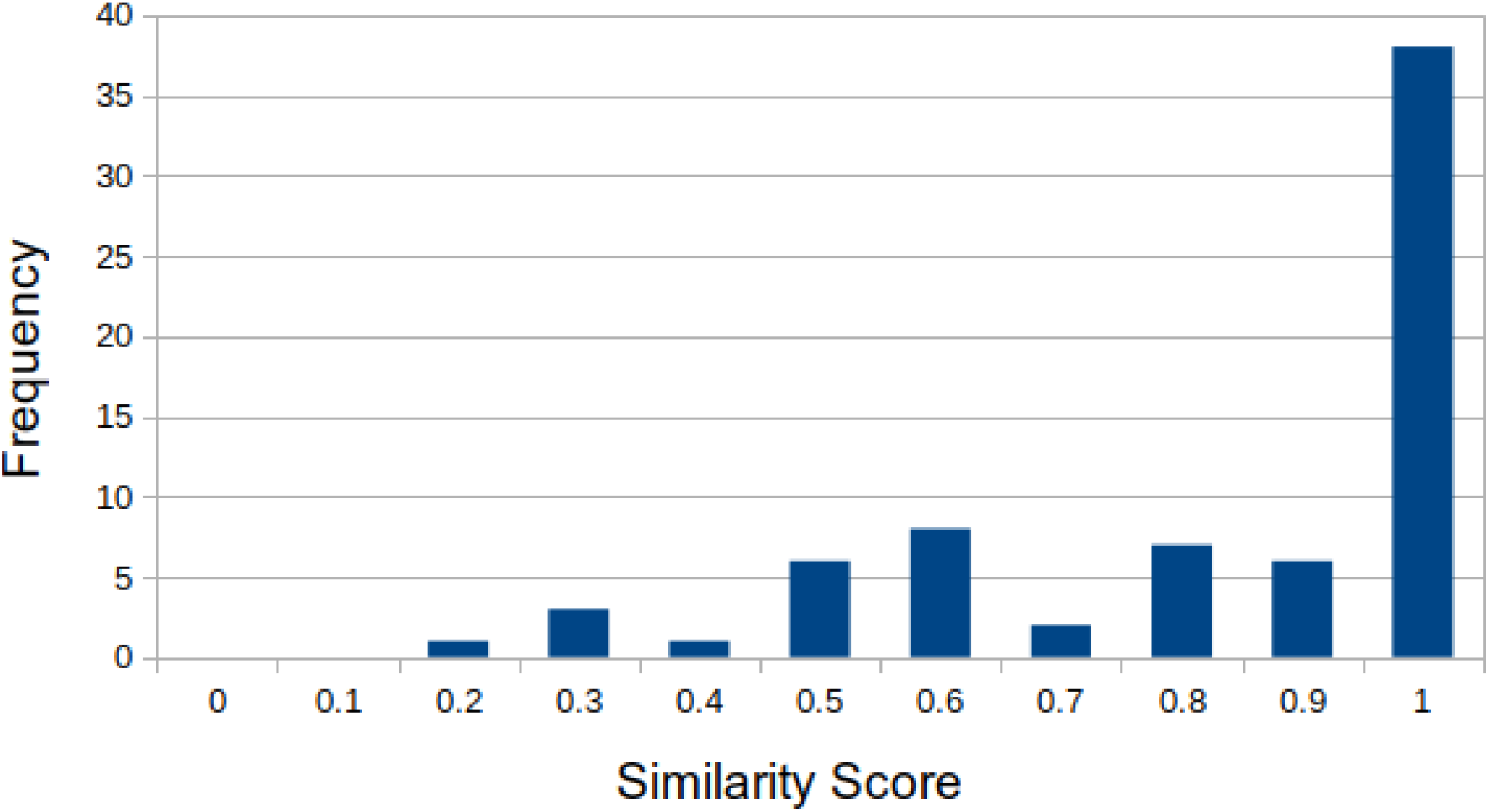
Distribution of motif similarity score in ChIP-seq dataset

##### B2) Comparison with DBD2BS

For comparison of DNA-sequence prediction we used DBD2BS, which generates a PWM for a given protein structure based on knowledge-based potentials (97). The 7 proteins that they have used are in their unbound conformation with an average RMSD of 1.1 Å. We have an average similarity score of 0.51 on this dataset. Since JEDII outputs a single DNA sequence and DBD2BS outputs a PWM, we could not compare the two methods directly. Therefore, we used their definition and compared the number of times the highest occurring nucleotide in the PWM is correctly predicted (Table 7). JEDII has a better performance than DBD2BS in 4 cases. In 1 case the number of correctly matched positions is the same for both methods, while in 2 of the 7 cases, DBD2BS performs better than JEDII.

##### B3) Comparison with the Farrel and Guo method

Though DBD2BS is the only algorithm that predicts the binding site for a given protein, there are other algorithms, which are specific for transcription factors. One such algorithm was developed by Farrel and Guo (96). They also predict PWMs from a protein structure, however instead of a template-based algorithm, they use a fragment-based method to generate a complex of a transcription factor with DNA. The sequence of DNA is then generated using permutations and combinations and evaluating each sequence with an energy function to output a PWM.

They evaluate the number of positions in the PWM that they have correctly predicted. We used this measure to compare the method with JEDII on the 27 protein chains they have used in their study. JEDII achieved an average similarity score of 0.62, while the Farrel and Guo method had a similarity score of 0.53. We performed better than Farrel and Guo in 18 out of 27 cases. In two cases, both methods had the same similarity score and in 7 cases the Farrel and Guo method had a better similarity score than JEDII (Figure 5).

**Figure 5:**
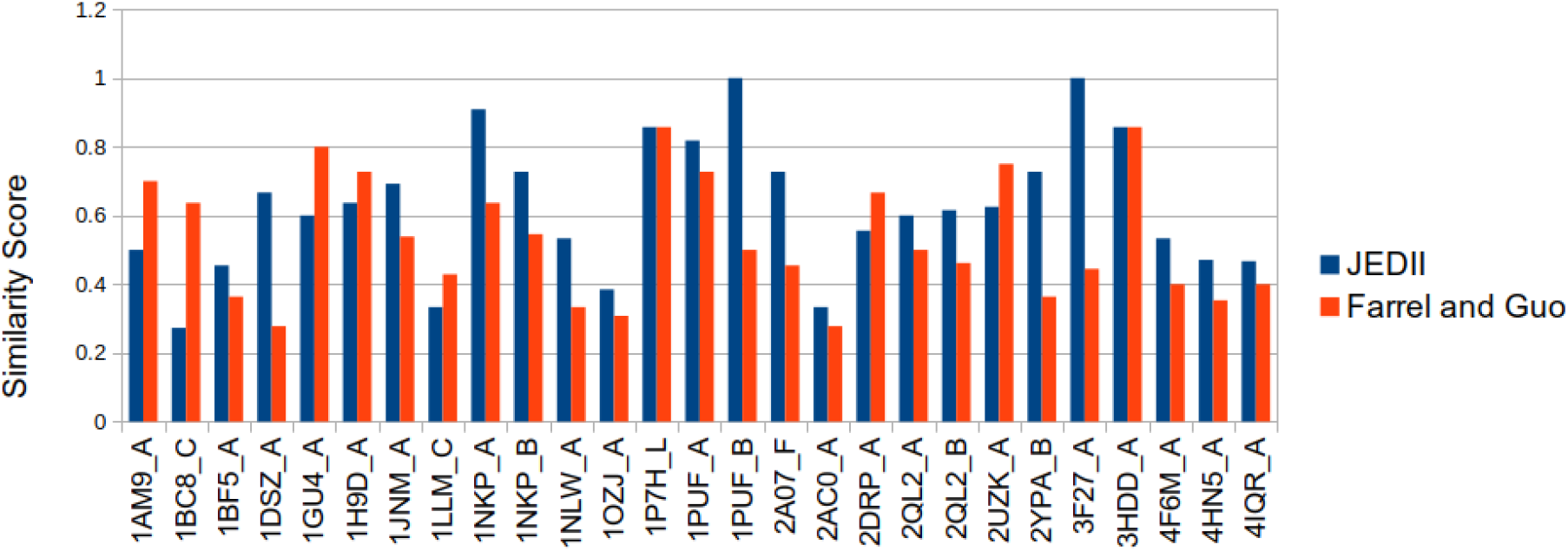
Comparison of JEDII with Farrel and Guo’s study on 27 transcription factor chains. The y-axis shows the similarity score for each chain mentioned on the x-axis. The similarity scores for JEDII and the Farrel and Guo method are depicted in blue and orange, respectively. The Farrel and Guo method had three algorithms for scoring. The best prediction for each protein chain was considered.

## Discussion

This study proposes an algorithm to predict the DNA sequence that would specifically bind a given protein structure. The method involves the superimposition of hydrogen bond donors and acceptors in the query protein against a dataset of known DNA-protein complexes. Hydrogen bonds are the most significant contributors to DNA-protein specificity (114–116). Therefore, we hypothesized the conservation of the orientation of hydrogen bond donors and acceptors in different DNA-binding proteins. Hence, the algorithm relies on the maximum match between the hydrogen bond donors and acceptors of the query and the template for template selection. We know that the protein region at the interface is complementary to the DNA. Therefore, if the query protein has matched well with the template protein, the DNA sequence of the template would also match with that of the query. Thus, we predict the DNA sequence of the template as the putative DNA sequence of the query.

We curated the PDNA-285 dataset to obtain only the protein-DNA complexes that exhibit sequence-specific binding. We used this dataset to test our hypothesis that the orientation of hydrogen bond-donor acceptor atoms shows conservation in specific DNA-protein complexes. The performance of JEDII is highest on the PDNA-285 dataset, lending credence to our hypothesis.

The MCC distribution of JEDII on the PDNA-285 dataset has a bimodal shape with peaks near 0 and 0.9. The sensitivity value is reflective of this distribution pattern. If we identify the optimal template, JEDII can predict the interface very accurately, while the binding site is completely missed in the other cases. We could identify the DNA sequences with an average similarity score of 0.62 on the PDNA-285 dataset. While we could identify the DNA-binding residues in 56 proteins with an MCC of 1, we obtained a similarity score of 1 for the DNA sequence prediction in 62 proteins. This implies that even though the DNA-binding pose may differ slightly, affecting the prediction of DNA-binding residues, our method can identify the DNA-protein interface correctly.

We analysed the contribution of the query identity and the average sequence identity between the template and query towards the MCC. Spearman’s correlation Coefficient was high for both parameters. However, the correlation with the query overlap was comparatively higher. Thus, JEDII tends to perform better in the presence of homologous structures bound to DNA. However, the performance is more dependent on the accuracy of the alignment between the query and the template proteins. To understand the contributions of the sequence identity between template-query, we employed the 40% NR template database to make predictions on the proteins from PDNA-285 dataset.

With the 40% NR template dataset, there was a reduction in MCC from 0.70 to 0.45. However, there were still 4 query proteins with an MCC of 1. The correlation between the MCC and the template-query sequence identity reduced to −0.05 from 0.60 with the 40% NR template dataset. However, the correlation between the MCC and the query identity remained high at 0.67. Therefore, in the absence of close homologs, the identification of DNA-binding residues relies on the structural similarity of the DNA-binding interface. There was a reduction in the efficacy of identification of the DNA sequence with the 40% NR template dataset. Since JEDII is a template-based method, a reduction in efficacy in the absence of close homologs was an expected outcome.

However, unlike other methods, there are examples where JEDII has a good performance without close sequential templates. One such example is that of the cro repressor. The selected template has a sequence identity of 31%, and the query and template have different CATH folds. Despite these differences, JEDII predicts the DNA-binding residues, DNA-binding pose and the predicted DNA sequence of the cro repressor protein with high accuracy. However, making uniformly good predictions in the absence of close homologs, needs improvements on the current algorithm. We plan to incorporate nucleotide changes based on the nucleotide-amino acid specificity rather than just relying on the geometric similarity of the interface.

Another evaluation of robustness was to check the performance of JEDII on unbound DNA-binding protein conformations. Previous studies have noted a conformational change in both DNA and proteins when they interact to form a complex (117, 118). Most of the studies utilizing structural information to predict DNA-binding residues look at the conformation of the protein in the DNA-bound state.

However, the aim of developing these methods is for the DNA-binding residue predictions of proteins that do not have a DNA-bound conformation. Thus, it would influence the accuracy of the methods in cases where conformations differ in the bound and unbound states. We used the APO-HOLO 82 dataset to analyze the performance of JEDII. While the accuracy does not change significantly with either apo or holo-structures as inputs, the MCC values are lower for the apo structure (0.55) compared to the holo-structure (0.67). Despite the conformational change, JEDII can identify 17/82 or 20% of the holo-structures as the templates for their respective apo structures, which points to its robustness.

The DNA sequence prediction is also unaffected with apo or holo-structures as inputs, with both giving a similarity score of 0.66. The identification of DNA sequence may have an advantage with the apo structure. JEDII identified the exact DNA sequence in 23 out of 82 cases using the holo-structure as query, while this number increased to 30 with the apo structure as input. The overall similarity score is the same for apo and holo, indicating that the selected templates for the holo structure have more near-native identifications than the apo protein.

We speculate that the template database does not have many structures similar to apo conformations. Thus if JEDII identifies the appropriate template, the DNA sequence matches perfectly. Otherwise, the similarity score is low. When we compare the performance of JEDII with other methods on this dataset, we outperform the existing methods on both datasets.

We can also obtain the unbound conformation of a DNA-binding protein through protein models. Especially with the development of AlphaFold2, there has been improvement in obtaining near-native conformations of proteins (104). Therefore, we tested the robustness of JEDII on models. While JEDII has a worse performance on both MODELLER and AlphaFold2 models than experimentally determined structures, the performance is slightly better with AlphFold2 models. This could be because AlphaFold2 models have more models with TM-scores >0.5 and also better average TM-score. Moreover, the RMSD between hydrogen bond donor and acceptor atoms, which are used for superimposition, is also lower for AlphaFold2 models.

GraphBind has a better performance with respect to MCC, sensitivity and F1 score when compared to JEDII while making predictions with protein models. This holds true for experimentally determined structures of the PDNA-129 dataset and models from this dataset. The PDNA-129 dataset contains 27 proteins whose structures have been determined by electron microscopy and the average resolution of these structures is 4.4Å. Since JEDII relies on finding appropriate templates using hydrogen bond-donor acceptor superimposition, the comparatively poor resolution may hinder the prediction accuracy. The template dataset, which is curated from X-ray crystal structures may also not have structures that resemble the proteins solved by electron microscopy. Therefore, future improvements to JEDII could involve incorporation of evolutionary information to overcome the lack of structural data.

While JEDII is a valuable tool, we wanted to compare its performance with the existing methods. Since no single method can predict both the DNA-binding residues and the DNA sequence, we compared the two kinds of predictions separately. For the DNA-binding residue prediction comparison, we used PDNA-62, P224 and PDNA-129 benchmark datasets. We perform better than other methods on the PDNA-62 and P224 datasets but not on the PDNA-129 dataset, on which GraphBind is better. We suspect GraphBind’s better performance has to do with their incorporation of evolutionary data. ~80% of its MCC contribution of GraphBind is from evolutionary information alone. As discussed before, the 27 electron microscopy structures also contribute to JEDII lagging behind GraphBind and incorporating evolutionary information is likely to improve our efficacy.

Apart from using benchmark datasets, we compared the performance of JEDII with GraphBind and HDock on the PDNA-285 dataset. GraphBind is currently the best method for predicting DNA-binding residues and incorporates both structural and sequential features for making predictions. While HDock is a protein-nucleic acid docking program that predicts the binding pose of the DNA. We perform better than GraphBind and HDock in all measures except the sensitivity.

To validate the DNA sequence prediction, we used data from ChIP-seq experiments, since it is one of the most common methods to identify the DNA sequence binding a protein. We used data from the PWMs generated by ChIP-seq experiments and compared the DNA sequence predicted by JEDII with all the sequences that would satisfy a given PWM. In this case, we have a motif similarity of 0.81.

One of the reasons for the high similarity score values in ChIP seq experiments could be the presence of ambiguous positions in the PWM. By ambiguous positions, we mean that all four nucleotide occur at that position (Ns) in the consensus sequence. While it is possible that at a particular position no single nucleotide is favored, the experimental conditions also influence the binding of proteins to DNA. Therefore, less stringent conditions could enhance non-specific binding resulting in a PWM with more Ns. Thus, it is crucial to rationalize the data obtained from ChIP-Seq experiments with known DNA-protein specificity patterns. Considering the time and resources involved in performing ChIP experiments, JEDII can be used to obtain a preliminary hypothesis of the putative binding site.

For comparing the DNA sequence prediction by JEDII with other computational methods, there were only two candidates-DBD2BS and a study by Farrel and Guo. While DBD2BS is a method that can be used for any DNA-binding protein, the method by Farrel and Guo is specific for transcription factors. DBD2BS aligns the query protein to templates and uses knowledge-based potentials to predict a PWM. Since JEDII can predict a single DNA sequence, thus for comparison, we used their measure of a correct prediction. They consider a prediction as correct if the predicted nucleotide is also the nucleotide with the highest frequency at that position. JEDII performs better than DBD2BS in 4 out of 7 test cases and equivalent in 1.

The Farrel and Guo method is a fragment-based method for predicting the PWM bound by transcription factors. Since a direct comparison was not possible, we compared JEDII with their method using similarity scores. For each position in the PWM, we consider it as a correct prediction if that nucleotide has the highest frequency for that position. The Farrel and Guo method also attempted to correctly identify a position in the PWM, but with the PCC method (119). We compared the similarity scores for both methods and performed better than the Farrel and Guo method in 18 out of 27 proteins. Both methods have the same similarity score in two cases, and JEDII has worse performance than the Farrel and Guo method in 7 cases. As mentioned previously, incorporating nucleotide-amino acid interaction probabilities could improve the performance of JEDII for DNA sequence prediction.

In the different datasets, a common limitation of JEDII is its comparatively lower sensitivity. The lower value of sensitivity is due to its inability to make a correct prediction without an appropriate template. With the increasing structural data, incorporation of evolutionary information and protein-nucleotide specificity information, JEDII can be improved in the future. Overall, despite being a solely structure-based method, JEDII outperforms not only other structure and sequence-based methods, but most methods that employ a combination of structure and sequence.

## Conclusion

We have developed an algorithm that can predict the DNA-binding residues, the binding pose and the DNA sequence that would bind a given protein structure. It is robust and can make predictions on the unbound conformation of proteins, in the absence of close homologs and on protein models. The unique feature of this method is that it can predict the DNA sequence for a given protein regardless of its structural or functional class. Thus, if sufficient structural data is available, this method could be extended to RNA-protein complexes as well. It integrates the problems of prediction of DNA-binding residues and the prediction of DNA sequences into the common umbrella of DNA-protein interface prediction.

## Acknowledgements

SN would like to acknowledge CSIR-Shyama Prasad Mukherjee Fellowship for funding.

