## Supplementary Table 1 for "JEDII: Juxtaposition Enabled DNA-binding Interface Identifier"

Supplementary Table 1: List of chains included in the PDNA-285 dataset

| Entry (PDB ID) | Chains |
|---|---|
| 1a0a | A,B |
| 1a3q | A,B |
| 1a73 | A,B |
| 1am9 | A,B |
| 1an4 | A,B |
| 1b3t | A,B |
| 1b94 | A,B |
| 1bdt | A,B,C,D |
| 1bf5 | A |
| 1bl0 | A |
| 1c9b | M |
| 1c9b | N |
| 1cf7 | A,B |
| 1cgp | A,B |
| 1cma | A,B |
| 1d02 | A,B |
| 1d3u | A,B |
| 1d66 | A,B |
| 1dc1 | A,B |
| 1dct | A |
| 1dh3 | A,C |
| 1dmu | A,B |
| 1dp7 | B,P |
| 1efa | A,B,C |
| 1f44 | A |
| 1f5t | A,B,C,D |
| 1fiu | A,B |
| 1fjl | A,B |
| 1fok | A |
| 1gd2 | E,F |
| 1gdt | A,B |
| 1gt0 | C,D |
| 1h9d | A |
| 1hcr | A |
| 1hlv | A |
| 1i3j | A |
| 1iaw | A |
| 1ic8 | A,B |
| 1if1 | A,B |
| 1io4 | A,B,C,D |
| 1jfi | A,B,C |
| 1jgg | A,B |
| 1jj4 | A,B |
| 1jt0 | A,B,C,D |
| 1k78 | A |
| 1k78 | B |
| 1kb6 | A,B |

| | |
|---|---|
| 1l3l | A,C |
| 1lli | A,B |
| 1lq1 | A,B |
| 1mdy | A,B |
| 1mnm | A,B,C,D |
| 1n3f | A,B |
| 1nkp | A,B |
| 1nlw | A,B |
| 1orn | A |
| 1ouz | A,B |
| 1owg | A,B |
| 1p7h | L,M |
| 1per | L,R |
| 1pp7 | U |
| 1pue | E |
| 1qps | A,B |
| 1r71 | A,B |
| 1r8d | A,B |
| 1rep | C |
| 1sax | A,B |
| 1skn | P |
| 1tro | A,C |
| 1u3e | M |
| 1u8b | A |
| 1vas | A |
| 1vkx | A,B |
| 1vol | A,B |
| 1xhu | A,B |
| 1yfi | A,B |
| 1yrn | A,B |
| 1ysa | C,D |
| 1z19 | A,B |
| 1z9c | A,B |
| 1zme | C,D |
| 1zs4 | A,B,C,D |
| 1zx4 | A,B |
| 2a66 | A |
| 2ady | A,B |
| 2bgw | A,B |
| 2bop | A,C |
| 2d5v | A |
| 2e1c | A |
| 2efw | A,B |
| 2er8 | A,B |
| 2evf | A |
| 2ex5 | A,B |
| 2ezv | A,B |
| 2fio | A,B |
| 2fqz | A,B |
| 2g1p | A,B |
| 2h27 | A,D |
| 2hap | C,D |

| | |
|---|---|
| 2i9k | A |
| 2ief | A,B |
| 2ih4 | A |
| 2itl | A |
| 2o49 | A |
| 2oq4 | A |
| 2p5l | C,D |
| 2pi0 | A,B,C,D |
| 2pue | A,C |
| 2pvi | A,B |
| 2qhb | A |
| 2ql2 | A,B |
| 2rbf | A,B |
| 2wt7 | A,B |
| 2ypb | A,B |
| 2yvh | A,B,C,D |
| 3bm3 | A,B |
| 3brf | A,D |
| 3bs1 | A |
| 3c25 | A,B |
| 3co7 | C |
| 3dw9 | A,B |
| 3e54 | A,B |
| 3fc3 | A,B |
| 3fd2 | A |
| 3fdq | A,B |
| 3fmt | A,B |
| 3g73 | A,B |
| 3g9i | A,B |
| 3gfi | A,C |
| 3gxq | A,B |
| 3gz6 | A,B |
| 3h0d | A,B |
| 3hqf | A |
| 3hqg | A,D |
| 3igm | A,B |
| 3ikt | A,B |
| 3jsp | A,B |
| 3jtg | A |
| 3jxb | C,D |
| 3kde | C |
| 3ket | A,D |
| 3ko2 | A,B |
| 3kov | I,J |
| 3lap | A,D |
| 3ldy | A,D |
| 3lsr | A,C |
| 3mkz | A,N |
| 3mln | A,B,E |
| 3mva | O |
| 3mx9 | A |
| 3mzh | A,B |

| | |
|---|---|
| 3ndh | A,B |
| 3nic | A,H |
| 3o9x | A,B |
| 3odh | A,B |
| 3on0 | A,B,C,D,E,F,G,H |
| 3oor | A |
| 3oqo | A,C,D,S |
| 3osf | A |
| 3pvv | A |
| 3q2y | A,C |
| 3q5f | A,B |
| 3qws | A,B |
| 3rmp | A,C |
| 3s8q | A,B |
| 3sqi | A |
| 3sse | A,B |
| 3tmm | A |
| 3u3w | A,B,P,Q |
| 3ukg | A |
| 3uvf | A |
| 3v1z | A,E |
| 3veb | A,B |
| 3vek | C |
| 3vw4 | A |
| 3vwb | A |
| 3w6v | A |
| 3waz | A,B |
| 3wty | A |
| 3wty | C |
| 3zhm | A |
| 3zkc | A,B |
| 3zpl | A,B |
| 3zqc | A |
| 3zql | A,B |
| 3zvk | A,B,C,D,E,F,G,H |
| 4atk | A,B |
| 4bhk | A,B |
| 4cja | A |
| 4e0j | A |
| 4egz | A,B |
| 4euw | A |
| 4fth | A,B |
| 4h10 | A,B |
| 4hf1 | A,B |
| 4hly | A |
| 4hqe | A,B |
| 4hri | A,B |
| 4i2o | A,B |
| 4iht | A,B |
| 4iqr | A,B |

| | |
|---|---|
| 4ix7 | A,B |
| 4j19 | A,B |
| 4jbm | A,B |
| 4jcx | A,B |
| 4jl3 | A,B,C,D |
| 4kny | A,B |
| 4kyw | A |
| 4l5s | A |
| 4l62 | A,B |
| 4ldx | A,B |
| 4lmg | A,D |
| 4lq0 | A |
| 4m8b | R |
| 4mte | A,B,C,D |
| 4ncb | A |
| 4nhj | A,B |
| 4on0 | A,B |
| 4osk | A |
| 4qtk | A,B |
| 4r4e | A,B |
| 4rb2 | C,D |
| 4rdm | A |
| 4u0y | A,B,C,D |
| 4ux5 | A,B |
| 4uzb | A,B |
| 4wls | A,B |
| 4wuh | A,B |
| 4xr2 | A |
| 4xrm | A,B |
| 4z5d | A,B |
| 4zq9 | A,B,C |
| 4zsf | A,C |
| 5a74 | A,B |
| 5ak9 | A |
| 5bmz | A,B |
| 5d2q | A |
| 5d5x | B,E |
| 5d8c | A,B |
| 5dwb | A,B |
| 5dy0 | A,B |
| 5e67 | A |
| 5ed4 | A,B |
| 5ego | A,B |
| 5eyb | A |
| 5f7q | C,E,J,L |
| 5fd3 | B |
| 5fmp | A,B |
| 5gnj | A,B |
| 5gpc | A,B,C,D |
| 5gzb | A |
| 5h3r | A,B |
| 5hlg | A,B |

| | |
|---|---|
| 5hnh | A |
| 5hr4 | C,J |
| 5hso | A,B,C,D |
| 5ity | A |
| 5j2y | A |
| 5j3e | A,B |
| 5jjv | B |
| 5jlt | A |
| 5jub | A,B,C,D |
| 5k1y | A,B,C,D,E, F |
| 5k7z | A,B |
| 5kkq | A |
| 5l6l | A,B,C,D,E, F,G,H |
| 5lxu | A |
| 5mhj | A,B |
| 5mpf | A,B |
| 5odg | A |
| 5t01 | A,B |
| 5tgx | A |
| 5trd | A,B |
| 5vpe | A,B |
| 5vvk | A |
| 5vvk | E,F |
| 5wjq | D |
| 5wx9 | A |
| 5x11 | A,B |
| 5x5l | B |
| 5x6e | A,B |
| 5xvp | A,B,C,D,E, F |
| 5yi2 | A,B |
| 5yj3 | C,D |
| 5yx2 | A,B,C,D |
| 5z7i | A,B,C |
| 6b0r | A |
| 6cro | A |
| 6el8 | A |
